# Entorhinal Cortex Wolframin-1-expressing neurons propagate tau to CA1 neurons and impair hippocampal memory

**DOI:** 10.64898/2026.02.19.706636

**Authors:** Jean-Christophe Delpech, Srinidhi Venkatesan Kalavai, Dhruba Pathak, Arun Reddy Ravula, Heena V. Lad, Emma C. Hays, Nina Touch, Andrea Rau, Charlotte Madore, Maria Medalla, Jennifer I. Luebke, Seiko Ikezu, Tsuneya Ikezu

## Abstract

Tau pathology emerges early in Alzheimer’s disease within entorhinal cortex layer II (ECII) and reaches hippocampal CA1, but how this circuit-level spread translates into sex-dependent vulnerability remains unclear. Using a circuit-defined model in which P301L human tau is expressed selectively in Wolframin-1 (Wfs1^+^) ECII neurons and propagates to CA1, we found that the extent and proximal–distal distribution of tau-positive CA1 neurons were comparable in males and females. Despite similar propagation, females exhibited broad hippocampal-dependent cognitive impairment (working memory, object recognition, fear acquisition, trace associative memory, and contextual fear memory), whereas males showed a selective deficit in trace associative memory. Consistent with these behavioral outcomes, CA1 pyramidal neurons in tau-propagated females displayed reduced excitability (slower action potential kinetics, reduced firing during depolarizing steps) and reduced spontaneous excitatory postsynaptic current (EPSC) amplitude, while males showed subtler intrinsic changes with altered EPSC kinetics. Bulk RNA sequencing of entorhinal cortex and CA1 revealed robust immune pathway engagement after tau propagation, with males showing a stronger Th1/Th2 and neuroinflammatory signature and CTLA4-associated signaling changes, whereas females showed prominent complement–phagosome pathway enrichment and a female-specific increase in Clec7a^+^ microglia density in CA1. CD4^+^ T-cell infiltration into CA1 was detected in both sexes. Together, these results indicate that sex-specific neuroimmune programs, rather than differences in tau propagation load, shape CA1 electrophysiological dysfunction and the breadth of memory impairment following early entorhinal-to-hippocampal tau spread.

## Introduction

Alzheimer disease (AD) pathophysiology includes the progressive accumulation of abnormally phosphorylated tau (pTau) in the brain, as one of the main hallmarks of the pathology [1]. Levels of pTau correlates significantly with synaptic loss and cognitive impairments. As per Braak post-mortem staging, tau pathology first appears in the superficial layers of the transentorhinal cortex and entorhinal cortex layer II (EC II), then spreads to the hippocampal Cornu Ammonis 1 (CA1) region in the prodromal stage of AD (Braak stage I-II) [2]. Tau spread is thought to take place among synaptically connected brain regions, however, the pathway that connects ECII to CA1 involved in this early tau spreading remains elusive. ECII neurons are heterogeneous across species and present two different chemical and morphological phenotypes: reelin^+^ multipolar stellate neurons and calbindin^+^ or woframin^+^ pyramidal neurons [2–5]. The majority of tau propagation animal models have demonstrated tau transfer from ECII stellate neurons to dentate gyrus, recapitulating tau spread visible in the advanced stages of AD. We recently showed that Wolframin-1 (Wfs1) positive cells in ECII can establish synaptic contacts with CA1 pyramidal neurons in the stratum lacunosum of the hippocampus through temporoammonic pathway [6]. We then demonstrated that tau-expressing ECII neurons can propagate tau to CA1, mimicking the early stages of tau pathology in AD [6]. This discovery paves the way to a new area of research to fully uncover the neuronal mechanism involved in tau propagation in early stages of AD. Our initial findings showed that tau propagates to CA1 similarly between male and female and induce functional alterations of the EC-CA1 pathway, culminating into EC-dependent associative fear memory deficits in both sexes.

Here, we aimed to assess hippocampal-dependent memories, including context-based spontaneous recognition of novelty and contextual-fear responses, and determine the neurobiological mechanisms involved at the transcriptomic level for the tau-induced alteration of CA1 neuronal functions, taking into account the sex. Our results indicates that although the extent of tau propgation to CA1 region is similar between Tau female and male mice, Tau females show significant reduction in cognitive functions as determined by multiple behavioral paradigms and impairment in the excitability of CA1 pyramidal cells, whereas Tau male mice show subtle changes in these paradigms. Transcriptomic profiling of EC and CA1 brain subregions reveal significant innate and cellular immune response in these regions after tau propagation, with female mice show specific induction of complement and activated microglial response compared to males. These data demonstrate vulnerability in female mice in behavioral and cellular response to tau propagation, and posit the role of innate and cellular immune response for the molecular mechanism.

## Methods

### Animals

Male and female Wfs1-Cre (gift from Dr Tonegawa, Riken-MIT, center for neural circuit genetics at the picower institute for learning and memory, department of biology and department of brain and cognitive sciences, Massachusetts institute of technology (MIT), Cambridge, MA) and C57BL6/J mice (Jackson Laboratories, Farmington, CT, USA) mice were kept on a 12-h light/12-h dark schedule with access to water and chow ad libitum. Pregnant mothers were single-housed. Offspring were weaned at P21 and group-housed with 2-5 same-sex littermates per cage. The number, sex and age of animals for each experiment are listed in each figure legend. No statistical methods were used to predetermine sample size, and randomization of samples was done according to age and litter. All animal procedures followed the guidelines of the National Institutes of Health Guide for the Care and Use of Laboratory Animals and were approved by the Boston University Institutional Animal Care and Use Committee and by Institutional Animal Care and Use Committee of the Mayo Clinic Florida. All data analysis was performed by researchers blinded to both genotypes and treatment groups.

### Stereotaxic surgeries

Stereotaxic viral injections were all performed in accordance with Boston University Institutional Animal Care and Use guidelines and Mayo Clinic Institutional Animal Care and Use guidelines. Mice were anaesthetized using 1-2% isoflurane in a 95% O2 and 5% CO2 mixture. Viruses were injected by using a computer-controlled stereotaxic frame (Neurostar, Germany) equipped with a glass micropipette attached to a 10-μL Hamilton microsyringe through a microelectrode holder filled with mineral oil. A microsyringe pump and its controller were used to control the speed of injection. The needle was slowly lowered (0.1 mm/min) to the target site, injections were done at a rate of 0.1 µL/min and the needle remained at the target site for 10 minutes after the injection. For histology, unilateral viral delivery into the right MEC of Wfs1-Cre mice was placed at Bregma coordinates: AP −4.85, ML +3.45, DV −3.30. The 24 to 32-week-old Wfs1-Cre mice were injected with 700 nl of AAV6-Flex-P301Ltau (3.34×10^11^ GC/mL) or 300 nL of AAV6-Flex-tdTomato (9.74×10^10^ GC/mL) and mice were subsequently perfused.

For electrophysiological recording and behavior, bilateral viral delivery into the MEC of Wfs1-Cre was placed at Bregma coordinates: AP −4.85, ML +3.45, DV −3.30 and AP −4.85, ML −3.45, DV −3.30. The 24 to 32 week-old Wfs1-Cre mice were injected with 700 nL of AAV6-Flex-P301Ltau or 300 nL of AAV6-Flex-tdTomato.

### Behavioral Testing

All behavioral tests were performed in an empty testing room within the animal vivarium, during the light cycle between the hours of 9 am – 6 pm. For all tests, animals were habituated in the test room for at least 1 hour before the start of testing. A test-free period of 5-7 d was used between behavioral tests. The lighting, humidity, temperature, and ventilation were kept as constant as possible in the testing room. The experimenter was not present in the room during any of the tests. The experimenter performing behavior tests was blind to the treatment group. All behavioral analyses were performed in a blinded manner. Animal sample sizes are reported in the Figure legends.

#### Y-maze test

The Y-maze assesses working memory based on the innate preference of a mouse to alternate arms while exploring a new environment. Typically, mice prefer to explore a new arm of the maze rather than returning back to the one that was previously explored. The Y-maze apparatus consisted of three arms with dimensions 35 cm (length) × 7 cm (height) × 5 cm (width) (San Diego Instruments). Testing was always performed in the same room and at the same time to ensure environmental consistency as previously described [7]. In the training session, one arm was closed with a gate and a test mouse was placed at the end of one arm and allowed to explore the two opened arms freely through the maze during a 5-min session. Then, the mouse was removed, the closing gate removed and the maze cleaned using 20% ethanol. After a 30 min time inter-interval, the test mouse was placed back in the Y-maze and allowed to explore the entire maze freely during a 5-min test session. The total amount of time spent in each arm (starting-familiar-new) was tracked using Ethovision XT14 software, and time in familiar vs novel arm was used to calculate an index of recognition. The maze was cleaned with 20% ethanol after each mouse to minimize odor cues.

#### Novel object recognition

During the training phase, mice were introduced with two identical objects two times during 5 min, separated by a 5 min interval (two training sessions). Animals exploring less than 2 sec with objects during the second 5 minute training session were excluded from the analysis. 24 h later, during the testing day, mice were introduced first during 5 minutes with the same two familiar objects. After a 5 min inter-trial interval, one object was replaced with a new object in the same arena at the same location as the familiar object, and exploration of the novel object was recorded during 5 min. Time spent by the mouse exploring each object during all the training and test sessions was measured manually by a blinded experimenter. Total time exploring the familiar and novel object during the final test session and an index of recognition calculated using the time exploring the familiar and the novel object during the final test session were reported.

#### Fear Conditioning

Trace fear conditioning experiment was performed in the animal facility during the light cycle following the previously published method that reported the involvement of the Wfs1 temporoammonic pathway in the trace fear memory paradigm with minor modification [8].

Trace paradigm: Wfs1-Cre mice injected bilaterally with either AAV-Flex-tdTomato or AAV-Flex-P301Ltau were placed 4 weeks after the injections in the context “A” and allowed to explore for 240 seconds, at which point a 20-s tone (85 dB, 2000 Hz) was played, followed by a 20-s trace and then a 2-s, 0.75 mA foot-shock. This was repeated two more times starting at 402 and 564 s for a total time of 706 s. On day2 (24 h later), mice were placed in a context “B” and allowed to explore for 240 s, at which point the same tone as day1 was played for 60 s, followed by 180 s of no-tone (post-tone period). This was repeated two more times for a total time in the context “B” of 960 s. On day3 (24 h later), mice were placed back in context “A” and allowed to explore for 300 s. Freezing behavior was recorded during all the time spent in either context.

### Whole cell patch clamp recording

#### Preparation of Brain Slices for Recording and Filling

A total of at least three mice per group were used for electrophysiological recordings, and at least 10 cells were recorded per animal. No formal statistical testing was used to pre-determine cell recording or imaging sample size, but we used sufficient sample size to ensure adequate power based on our previous published studies [9] and those generally employed in the field. Mice were fully anesthetized with isoflurane before decapitation with guillotine. The brain was immediately removed and placed into ice-cold oxygenated Ringer’s solution (concentrations, in mM: 26 NaHCO_3_, 124 NaCl, 2 KCl, 3 KH_2_PO_4_, 10 Glucose, 2.5 CaCl, 1.3 MgCl_2_; pH 7.4, chemicals from Fluka). The right hemisphere was placed against a 3% agar block for support and sliced at 300-µm thickness with a Leica VT1000 S vibrating microtome (Leica Microsystems) yielding 4 coronal slices containing the dorsal hippocampus. Slices were maintained at 33.3 oC in oxygenated Ringer’s solution in a holding chamber for equilibration for at least 1 h prior to field potential recordings. Following this equilibration period, slices were transferred to submersion-type recording chambers at room temperature (Harvard Apparatus, Holliston, MA) affixed to the stages of Nikon E600 infrared-differential interference contrast (IR-DIC) microscopes (Micro Video Instruments, Avon, MA). Slices were visualized under IR-DIC optics using an Andor Xyla digital camera. Electrodes were created from borosilicate glass with a Flaming and Brown micropipette puller (Model P-87, Sutter Instruments). Pipettes were filled with a potassium methanesulfonate internal solution (concentrations, in mM): 100 potassium methanesulfonate, 15 KCl, 3 MgCl_2_, 5 EGTA, 10 Na-HEPES, 1% biocytin (1% n-biotinyl-L-lysine), and had a resistance of 4-7 MΩ in external Ringer’s solution. Standard whole cell patch clamp recording techniques were used to examine the electrophysiological properties of the cells.

#### Physiological Inclusion Criteria

Neurons were recorded from prefrontal cortical areas including the prelimbic area, anterior cingulate, and dorsal premotor areas, by experimenters blind to the treatment groups. Cells were visually identified as CA1 neurons and were required to have stable access, low noise, and a resting membrane potential below −55 mV for inclusion in the study.

#### Physiological Analysis

PatchMaster acquisition software and EPC-9 or EPC-10 patch-clamp amplifiers (HEKA Elektronik) were used to acquire electrophysiological data including passive membrane, single action potential, and repetitive firing properties. A total of 11 physiological characteristics were assessed by experimenters blind to the treatment groups. FitMaster Analysis Software (HEKA Elektronik) was used to analyze passive and active membrane properties. The passive membrane properties, resting membrane potential (Vr), input resistance (Rn), and the membrane time constants (Tau) were recorded under current clamp conditions. Vr was measured as the voltage present when the current injection was zero. Rn and Tau were measured by injecting 20 mV steps starting at −160 mV and ending at +100 mV. Tau was defined by fitting the membrane response to a small current injection (−20 or −40 mV) to a single exponential function. Rn was assessed as the slope of a best-fit line through the V-I plot. Single spike properties [threshold, amplitude (amp), rise, decay, and duration at ½ amplitude (Dur ½)] were measured from the first single spike recorded on the 200 ms current step. Threshold was measured at the point the action potential’s slope was greater than 1 mV/ms. The amplitude was the maximum voltage recorded at the peak of the spike. Single spike rise was the time, in ms, for the action potential to rise from threshold to peak amplitude. Fall time was the time for the action potential to return to threshold from the peak amplitude. Dur ½ is the duration of the spike from half amplitude on the rise to half amplitude on the fall of the action potential. Repetitive firing of APs was determined as the number of spikes evoked at each step in a series of current steps that increased by 20 mV from −100 to +120 mV. sEPSCs were recorded for 2 min at a holding potential of −80 mV. sIPSCs were recorded at a holding potential of −40 mV. Following acquisition of spontaneous synaptic data, tetrodotoxin (1 µM) was added to the Ringer’s perfusion medium and, following a 5 min equilibration period, action potential independent mEPSCs were recorded as above. Synaptic data was assessed using MiniAnalysis software (Synaptosoft), with event detection threshold set at maximum RMS noise level (5 pA). The sEPSCs and sIPSCs were automatically detected and manually confirmed to yield frequency and mean amplitude data. Averaged waveforms were created from all the EPSCs or IPSCs acquired for each neuron. From this, we calculated average EPSC and IPSC rise and decay time constants, and area under the curve (pA/ms).

### RNA sequencing

Brains were collected after cold PBS perfusion, and PFC and HPC were dissected on ice. Brain regions were immediately homogenized using pestle tissue grinder in Qiazol before freezing the samples and storage at −80°C. Total RNA was then extracted using the miRNeasy Micro Kit (Qiagen 217084). RNA concentration and purity were verified with a Bioanalyzer 2100 (Agilent Technologies), and all samples had a RIN score above 9.0. RNA libraries were prepared using 200 ng of total RNA following the Illumina Stranded mRNA Ligation Sample Prep Kit protocol. Library concentration and size distribution were assessed using a Bioanalyzer DNA 1000 chip (Agilent Technologies) and Qubit fluorometry (Invitrogen). Libraries were sequenced to obtain 50 million fragment reads per sample, adhering to Illumina’s standard protocol on the NovaSeq™ 6000 SP flow cell. Sequencing was performed as 100 × 2 paired-end reads using the NovaSeq SP sequencing kit and NovaSeq Control Software (version 1.8.0). RNA quality checks and RNA-seq were conducted by the University of Chicago Genomics Core, yielding 30 million paired-end reads via the Illumina NovaSeq 6000, using the oligo dT directional method for library preparation.

Reads were mapped to the mouse reference transcriptome GRCm39, version M27, using STAR (version 2.6.1d). Quality checks were performed with FastQC and MultiQC. Reads were trimmed with trimmomatic package, followed by feature count generation with Rsubread package. Filtering was performed to eliminate non-protein coding genes and genes with low expression across multiple samples (i.e., normalized counts < 5 in more than half of the samples for each sex in for each treatment group). Global differential analyses were carried out for each brain region using DESeq2 version 1.28.1, with fixed effects to account for batch as well as sex and treatment group (Saline+CTRL; MIA+CTRL; Saline+REP; MIA+REP). Analogous differential analyses were also performed for each sex independently, with fixed effects for batch and treatment group. Treatment-specific differentially expressed genes (DEG) between each pair of treatment groups were identified using contrasts and a significance threshold of 10% for the false discovery rate (FDR) K-means clustering was used to cluster normalized expression Z-scores of DEGs to identify modules of distinct transcriptional signatures. Pathway enrichment analysis was performed using clusterProfiler version 4.10.1and Ingenuity Pathway Analysis® software. Upstream regulator analysis was done using Ingenuity Pathway Analysis® software. Data wrangling and graphic generation were conducted using the tidyverse suite of R packages (R version 4.0.0).

### Statistical analysis

Data are presented as means ± standard error of the mean (s.e.m.). Comparisons between groups were done by Student’s *t*-tests. Multiple comparisons were performed by either one-way or repeated measures ANOVA, followed by a Student–Newman–Keuls’s *post hoc* test after assessing for data normality. Data analyses were performed using Prism 10.0 (GraphPad). Outliers could be identified and removed using Rout test and Q = 1%. A statistically significant difference was assumed at p <0.05.

### Immunohistochemistry

Brains were collected after perfusion and fixation with 4% PFA in PBS, followed by overnight post-fixation and cryoprotection in 30% sucrose and embedding in an OCT compound (Sakura). Brains were frozen on dry ice and sectioned by cryostat at 30-µm thickness into sagittal sections. For the lymphocyte IHC and microglial Clec7a studies, three sections collected at every 360-μm interval were used for IHC (ML 2.88 - 2.16 mm). We also confirmed first tau propagation using human tau antibody staining. For tau propagation analysis in sub-regions of the CA1, one section every 150 µm was used for immunohistochemistry.

Sections were washed with PBS for 10 min prior to antigen retrieval in 10 mM tris base/1 mM EDTA/PBS at 95°C or 10 mM sodium citrate at 80 C° for 20 minutes followed by 20 min at room temperature (RT). Sections were then washed three times for 5 minutes in PBS then blocked in 5% normal goat serum, 5% bovine serum albumin (BSA), and 0.3% Triton-X in PBS for 1 hour at RT. Primary antibodies diluted with 5% BSA/0.1% Triton-X in PBS and sections were incubated for 24hrs at 4°C. The following antibodies and reagents were used: mouse anti-human tau (1:1000; ThermoFisher Scientific, HT7, MN1000B), mouse anti-CD4 (1:500, BD,553043), rat-anti-CD8 (1:500, ThermoFisher Scientific, 14-0808-82), rat anti-Clec7a (1:300, Invivogen, Mabg-mdect), rabbit anti-WFS1 (Proteintech, 26995-1-AP; 1:1000). Sections were then washed three times in PBS and incubated with Alexa Fluor 488–, Alexa Fluor 594–, Alexa Fluor 568–, Alexa Fluor 647–conjugated antibodies or streptavidin–Alexa Fluor 488–conjugated antibody (all from Invitrogen; 1:1000) for 2 h at room temperature. Following staining, sections were washed with PBS with 0.02% Triton-X and then mounted using Fluor mount-G with DAPI (Invitrogen, 00-4959-52).

For IHC analysis, we used FIJI software. We analyzed CA1 in brain sections of 30 µm every 150 µm from bregma 3.2 to 0.9 mm. We fist took large images of each brain section and used DAPI and WFS1 signal to identify the exact localization of the section compared to Allen Atlas for bregma coordinate. CA1 and DG were then identified based on DAPI staining, clearly allowing us to distinct CA1 from CA2 and subiculum and identify DG, knowing that WFS1 is only expressed in CA1 pyramidal neurons. CA1 and DG were manually cropped and areas measured on each image. In order to determine HT7^+^ neurons in proximal and distal CA1 areas, we quantified the total number of HT7^+^ neurons in sub-regions of CA1 in each image to be able to measure the propagation in these brain sub-regions. To do so, we manually defined for each image the value of intensity needed to count a neuron as positive. This value was at least two times superior to the background signal measured on a DAPI^+^ and HT7^−^ neuron of the corresponding image. We counted the total number of HT7^+^ neurons for CA1 of each image and then calculated the density of HT7^+^ neurons for each image by dividing the the total number of HT7^+^ neurons in CA1 by CA1 areas. Final results were presented for each animal quantified (N = 4 total) from each brain section per animal per sex.

For IHC analysis of CD4^+^, CD8^+^ and Clec7a^+^ cells, we used FIJI software for the analysis of CA1 region using 30- μm thickness brain sections in every 360 μm internval (three sections/animal) from ML 2.8 to 2.16 mm. We first took large images of each brain section and used DAPI to identify the exact localization of the section and region of interest compared to Allen Atlas for bregma coordinate. To quantify positive cells, we manually defined for each image the value of intensity needed to count a neuron as positive. This value was at least two times higher than the background signal measured on a DAPI^+^ and CD4^−^, CD8^−^ and Clec7a^−^ cell of the corresponding image. We counted the total number of CD4^+^, CD8^+^ and Clec7a^+^ cells for each ROI of each image and then divided it by the ROI’s area.

### Imaging

Large field fluorescence images of the mouse brain sections were acquired using an epifluorescence microscope (Nikon Eclipse Ti, Japan), using Plan Fluor ELWD 20× Ph1 DM objective at a 1024×1024-pixel resolution at excitation wavelengths of 405, 488, 561, and 647 nm.

## Results

To induce tau propagation from ECII Wfs1^+^ neurons to CA1 pyramidal neurons, we employed Wfs1-Cre transgenic mice expressing Cre recombinase under the Wfs1 promotor [6, 8], and injected biliteraly Cre-inducible AAV expressing human 2N4R P301L Tau mutant (AAV2/6-Flex-P301Ltau) into the MEC of Wfs1-Cre mice (**Fig.1a**). At 4 weeks post-infection, we confirmed our previous data, showing a restricted and strong HT7 signal in ECII Wfs1^+^ neurons (**Fig. 1b-d**) as well as HT7^+^ cells in the subiculum and the CA1 including the sl and the pyramidal layer but not in the DG (**Fig. 1c-e**) of males and females mice [6]. We previously reported no difference between males and females in terms of the total number of HT7 positive CA1 neurons [6]. CA1 actually present functional heterogeneity across anatomical axes, with the transverse axis indicating spatial encoding being more robust towards CA2 (i.e. proximal CA1) and non-spatial encoding being more robust towards subiculum (i.e. distal CA1) [10, 11]. We thus evaluated the amounf of HT7 positive CA1 neurons according to transverse axis localization and found a comparable HT7 positivity in proximal and distal CA1 region in both sexes (**Fig. 1f**). The data suggest that there is no sex-dependency in the tau pathology development.

**Fig. 1.**
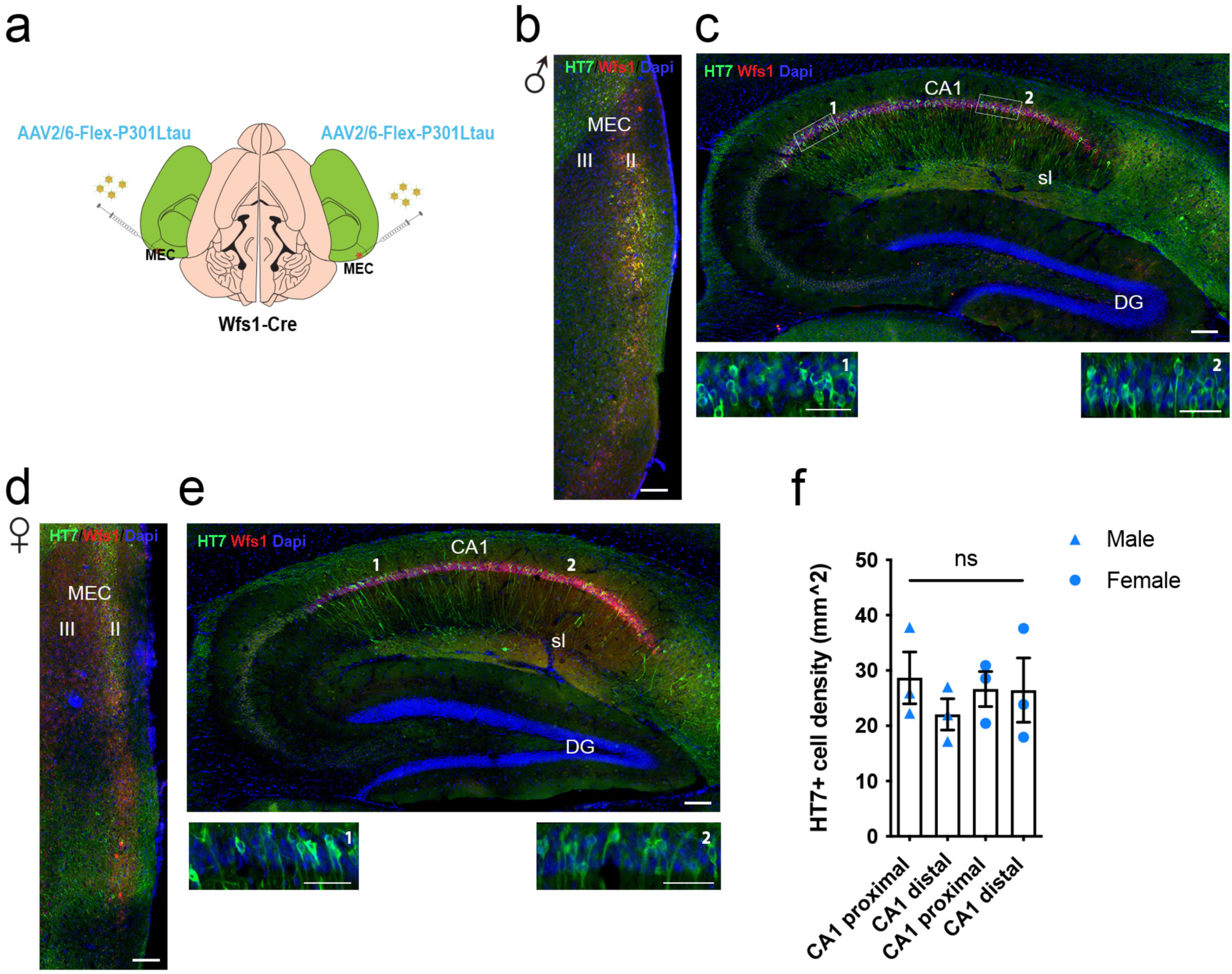
Wfs1^+^ neurons in entorhinal cortex upper layer II project and propagate human tau to distal and proximal CA1 in both male and female. **(a)** Diagram of the mouse brain indicating the location of AAV6-Flex-P301L human tau bilateral injections into the MEC of Wfs1-Cre mice. **(b-e)** Parasagittal section of the MEC and HPC of Wfs1-Cre mouse injected with AAV6-Flex-P301Ltau immunostained with anti-human tau HT7 (green) and anti-Wfs1 (red) 4 weeks after MEC injection in males (**b-c**) and females (**d-e**). **(f)** Quantification of HT7^+^ neurons in proximal and distal CA1 of Wfs1-Cre mice injected with AAV6-Flex-P301Ltau immunostained with anti-human tau HT7 at 4 weeks after MEC injection (Average: two-way Anova F (1, 8) = 0,62, ns=non significant, N=3 animals). Graph indicates mean±s.e.m. Scale bars = 100 µm.

The hippocampus is a brain structure necessary for spatial memory acquisition [12, 13], which are one of the first memory affected in AD [14]. We thus evaluated behavioral phenotypes using a series of tests to assess memory abilities known to depend upon hippocampal integrity, including contextual novelty recognition as a measure of spatial working memory and fear-dependent contextual end temporal memories as a measure of spatial associative memory abilities in both sexes 4 weeks after Tau EC injection (**Fig. 2a**). We aim to compare Wfs1-cre mice injected bilaterally in the ECII either with an AAV expressing tdTomato (AAV2/6-Flex-tdTomato), defined as Control, and with an AAV expressing human 2N4R P301L Tau mutant (AAV2/6-Flex-P301Ltau), defined as Tau. We tested them in the following order: Y-maze forced alternation, novel object recognition and fear conditioning tests (**Fig. 2b**). We first examined if tau interferes with spatial working memory and used the forced alternation Y-maze test (**Fig. 2c**). Control male mice spent significantly more time exploring the novel arm compared to the familiar arm, similarly to Tau male mice (**Fig. 2d**), allowing them to present a novelty discrimination index significantly above chance level (**Fig. 2e**), indicating a normal spatial working memory ability in male mice. In contrast, control female mice spent more time exploring the novel arm compared to the familiar arm, whereas Tau female mice spent the same amount of time exploring both arms (**Fig. 2f**). Control female mice only presented a novelty discrimination index significantly above the chance level (**Fig. 2g**), indicating that Tau propagation into CA1 affects spatial working memory ability in female mice. All the animals explored the three arms during the test. When exploring total travelled distance or, average velocity, we found no difference in male mice (**Suppl. Fig. 1a**) and a significantly increased distance and velocity in Tau female mice compared to control group (**Suppl. Fig. 1b**). To gain confidence into the Tau effect onto working memory abilities, we then conducted a novel object recognition test, which is mainly regulated by the perirhinal cortex and hippocampus (**Fig. 2h**) [15]. We confirmed that control and Tau male mice present normal working memory abilities in this test. Indeed, control and Tau male mice spent significantly more time exploring the novel object compared to the familiar one, reflected in a significant recognition index above the chance level (**Fig. 2i-j**). In regards to female mice, we also confirmed that Tau females present a deficit in working memory abilities. Only control female mice spent significantly more time exploring the novel object compared to the familiar one, as reflected in their significant recognition index above the chance level (**Fig. 2 k-l**). When looking at the total exploration time or frequency of interaction with the two objects during the training phase, we could not find any difference between the control and Tau groups in either sexes (**Suppl. Fig. 2.a-b**). Taking the two tests assessing working memory abilities into account, we found that only female Tau mice present a deficit. Finally, we examined whether Tau spread from the ECII to CA1 could affects episodic memory in a sex dependent manner, consisting in the association of objects with time and space [16], and relying on the enthorinal cortex-hippocampus network in animals and humans [17]. The medial EC is engaged in the “general” information about space and interacts with the hippocampus in this function [18]. In addition, the temporoammonic pathway between the EC and CA1 is involved in acquisition and reinstituting of trace memories where associated stimuli are presented with a delay to the animals [8]. Indeed, trace memories, defined as the temporal association of discontinued events, are driven by direct inputs from the ECIII neurons projecting to the CA1 pyramidal neurons [19] and modulated by the ECII neurons projecting to the CA1 ^7^. The CA1 pyramidal neurons are also directly involved in the formation and retrieval of episodic memories, including contextual memories [12, 13]. Thus, we used the trace fear conditioning to test trace and contextual memories in control and Tau mice in both sexes. When we looked at the acquisition phase, control and Tau groups significantly increased their freezing time after the three sound-shock associations, with male mice behaving normally during the acquisition (**Fig. 2m; Suppl. Fig. 3a**), and Tau female mice presenting a significantly lower amount of time freezing across the acquisition phase (**Fig. 2n**) without significant differences visible when focusing on the specific baseline, tone and post-tone period of the acquisition (**Suppl. Fig. 3c**). During the test phase, Tau group showed an altered trace fear memory, as evidenced by an overall reduction in their freezing time in Tau females (**Fig. 2o,p**) and by a reduced freezing time during the tone presentations in Tau males (**Fig. 2q**), or during the post-tone times in Tau females (**Fig. 2r**). Additionally, only the Tau females showed a deficit in contextual fear memory, as can be seen from the similar freezing time in Tau males and the reduced freezing time in Tau females compared to their respective control groups (**Fig. 2s,t; Suppl.Fig. 3b,d**). Overal, we found that both male and female Tau mice display a deficit in trace and only Tau female present a deficit in contextual memories, reflecting a common temporoammonic pathway dysfunction and a female specific CA1 dysfunction. In summary, these behavioral tests demonstrate that Tau female mice show cognitive impairment in forced alternation, novel object recognition, fear memory aquisition, trace associative memory and contextual fear memory, whereas Tau male mice exclusively showed impairment in trace associative memory, demonstrating more susceptibility to tau pathology-induced cognitive impairments in females.

**Fig. 2.**
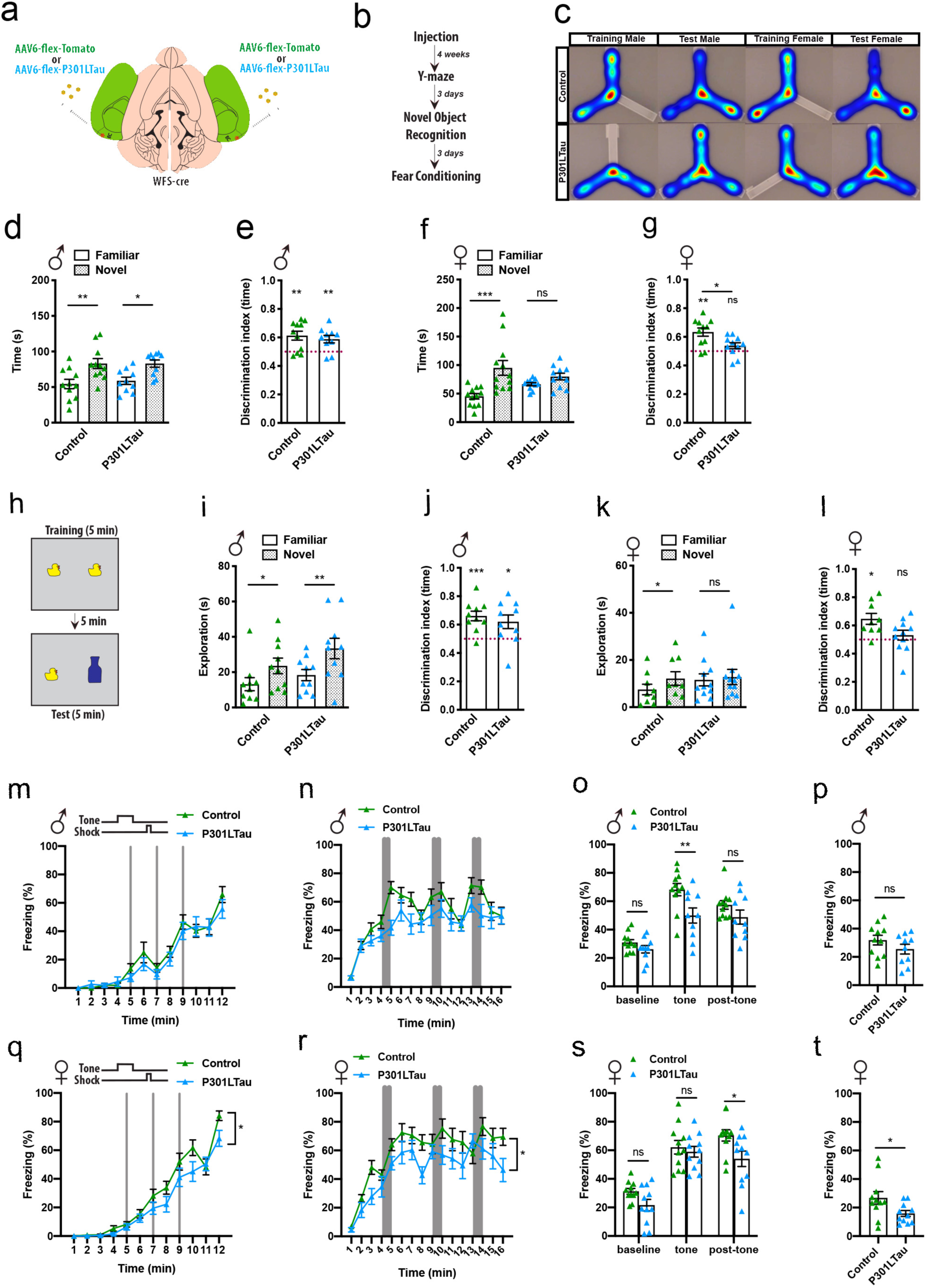
Tau propagation into CA1 induces more pronounced memory deficits in female. **(a)** Diagram of the mouse brain indicating the location of AAV6-Flex-P301L human tau or AAV6-Flex-tdTomato bilateral injections into the MEC of Wfs1-Cre mice. (**b**) Shematic representation of the behavioral tests conducted after bilateral cerebral injections of Wfs1-Cre mice. (**c**) Y-maze assay representative tracking images. (**d**) Duration of exploration of the Y-maze arms in male during the test session showing no difference between tomato and tau groups. (**e**) Ratio of the time exploring the novel arm over the novel and familiar arms in male showing no differences among the groups. (**f**) Duration of exploration of the Y-maze arms in female during the test session showing an absence of recognition of the novelty only in the tau group. (**g**) Ratio of the time exploring the novel arm over the novel and familiar arms in female showing an absence of recognition of the novelty only in the tau group. (**h**) Scheme of the novel object recognition paradigm showing the presence of a novel object during the test session. (**i**) Time exploring the familiar and novel objects showing novelty recognition among the groups in males. (**j**) Ratio of the time exploring the novel object over the novel and familiar objects in male showing novelty recognition among the groups. (**k**) Time exploring the familiar and novel objects showing novelty recognition only in tomato group in females. (**l**) Ratio of the time exploring the novel object over the novel and familiar objects in female showing novelty recognition only in tomato group. (**m**) Time-course of freezing observed in the fear conditioning paradigm during training on day 1 showing no differences among the groups in males. (**n**) Time-course of freezing observed during Trace-memory testing on day 2 in males showing overall no significant difference among the groups. (**o**) Average freezing time during baseline, tone and post-tone periods measured during trace-memory testing on day 2 in males, showing a significant reduction in freezing in tau group during tone periods. (**p**) Average freezing time during contextual-memory testing observed on day 3 in males, showing no difference among the groups. (**q**) Time-course of freezing observed in the fear conditioning paradigm during training on day 1 showing a significant reduction in females. (**r**) Time-course of freezing observed during Trace-memory testing on day 2 in females showing a significant reduction in tau group. (**s**) Average freezing time during baseline, tone and post-tone periods measured during trace-memory testing on day 2 in females, showing a significant reduction in freezing in tau group during post-tone periods. (**t**) Average freezing time during contextual-memory testing observed on day 3 in females, showing a significant reduction in tau group. Two-way ANOVA or two-way ANOVA on repeated measures, Tukey’s post-hoc, One-sample t-test for ratio (e,g,j,l), Unpaired-t-test (p,t), *<0.05, **< 0.01, ***< 0.001, Graph indicates mean±s.e.m.

**Fig. 3.**
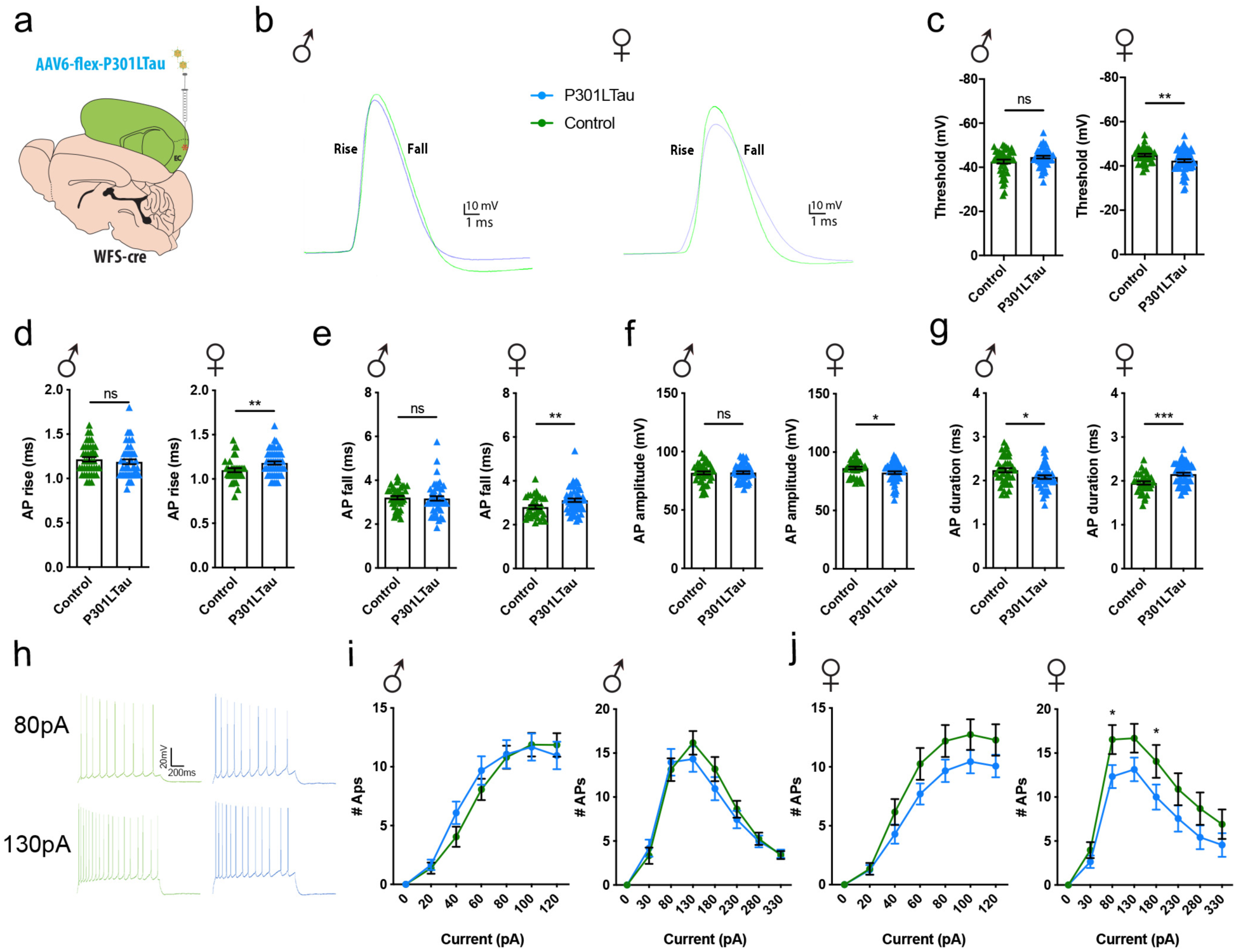
CA1 neuronal intrinsic properties are affected in female after Tau propagation. **(a)** Diagram of the mouse brain indicating the location of AAV6-Flex-P301L human tau or AAV6-Flex-tdTomato bilateral injections into the MEC of Wfs1-Cre mice. (**b**) Representative traces of action potentials from CA1 pyramidal neurons in slices prepared from control tomato and tau groups in both male (felt) and female (right). (**c-g**) Intrinsic properties of pyramidal neurons of CA1 in both male and female and tomato control and tau groups. (**c**) threshold. (**d**) AP rise. (**e**) AP fall. (**f**) AP amplitude. (**g**) AP duration. (**h**) Representative repetitive action potential firing responses to a +120pA current step of CA1 pyramidal neurons from control tomato and tau groups. (**i-j**) Number of action potential triggered by current steps applied to brain slices prepared from control tomato and tau groups in both males (**i**) and females (**j**). Two-way ANOVA on repeated measures, Tukey’s post-hoc (i-j), Unpaired-t-test (c,-g), *<0.05, **< 0.01, Graph indicates mean±s.e.m.

We previously reported an altered excitability of CA1 neurons after Tau propagation in ECII-CA1 Tau mice [6]. From the differences we found in hippocampal-dependent memories in the current study, we decided to examine CA1 neuronal properties, aiming to better understand the neurobiological substrate at the origin of the behavioral effects we found specific to females. The whole cell patch-clamp recording of CA1 pyramidal neurons demonstrated that most of the changes in neuronal intrinsic firing properties were seen in Tau females instead of Tau males at 4 weeks post surgery (**Fig. 3a-b**). Tau female mice show significant reduction in the action potential (AP) threshold (**Fig. 3c**), significant increase in AP rise, fall and duration and an increase in AP amplitude compared to controls (**Fig. 3d-g**). Conversely, Tau males only show significant reduction in AP duration compared to controls. There were no effects of Tau on the other intrinsic properties in both sexes (**Suppl. Fig. 4a-f**). These data indicate a significant reduction in neuronal excitability of CA1 pyramidal cells in Tau females specific manner. When we applied current steps to elicit AP, we found no differences between control and Tau males, but we did observe a significant reduction in the number of AP when more than 80 pA of current was applied to Tau females when compared to controls (**Fig. 3h-j**), confirming reduced excitability of CA1 pryamidal cells in Tau females. We next measured spontaneous excitatory postsynaptic currents (sEPSCs) and spontaneous inhibitory postsynaptic currents (sIPSCs) in CA1 pyramidal cells. We found only a significant reduction in amplitude in Tau female compared to control, and a significant reduction in rise and decay in Tau male compared to controls (**Suppl. Fig. 5**). Parameters of inhibitory postsynaptic currents showed no specific effects in both sexes. In summary, CA1 pyramidal neurons in Tau female group exhibit functionally significant changes in intrinsic properties, including slower AP kinetics, reduced number of AP elicited after current steps, and reduced amplitude in spontaneous excitatory postsynaptic currents, whereas Tau males had a significant reduction in AP duration with concurrent elevation and decay in spontaneous excitatory postsynaptic currents.

Neuronal proper functioning depends on spine morphology and density. Therefore, we additionally assessed the effect of Tau propagation on dendritic spine density in biocytin-filled and electrophysiologically characterized CA1 pyramidal neurons as previously described ^19^. We found no differences in dendritic spine density between Tau and control groups in both males and females (**Suppl. Fig. 6**). In order to disentangle the neurobiological mechanisms involved with the impact of Tau propagation from ECII to CA1, taking into account the sex, we sought a transcriptomic approach on brain regions dissected specifically to study CA1 and EC. Specifically we selected and dissected the prefrontal cortex (named PFC), as a negative control region we previously demonstrated had an absence of tau propagation [6], the enriched dorsal CA1 region of the hippocampus (named CA1, without DG, nor ventral hippocampus region), and the enthorinal cortex (named EC). We then performed bulk RNA sequencing from brain region homogenates of male and female Tau and tdTomato groups. Principal component analysis of the regions showed that male and female cluster discretely in the 3 regions. We also found that Tau and tdTomato samples cluster separately in EC and CA1, but not in PFC, corroborating previous findings of PFC as a less affected region and, substantiating the propagation of Tau in the EC and CA1 (**Suppl. Fig.7**). A heatmap plotted of the differentially expressed genes (DEGs) across each tissue region and experimental group (**Fig. 4A**), confirm that the expression profile of genes cluster accordingly. Specifically, a cluster of genes are upregulated with the Tau group compared witht the tdTomato group in both sexes, with a potentially more pronounced effect in the EC; a visible effect in CA1 due to Tau propagation. As expected, the expression profile across the PFC dissected from the P301L Tau group paralleled the expression profile of the PFC within the control group. Overall, with the EC and CA1, we found more DEGs in males (780 total DEGs) compared to females (230 total DEGs) (**Fig. 4b, Table1**) and an overall increased number of upregulated genes (**Fig. 4c**). We found that most of the DEGs in EC female group are shared with the other contrasts, except for 16 unique DEGs. Interestingly, there were 399 unique DEGs in the EC male contrast group, with an additional 381 shared DEGs with the other contrasts (**Fig. 4b**).

**Fig. 4.**
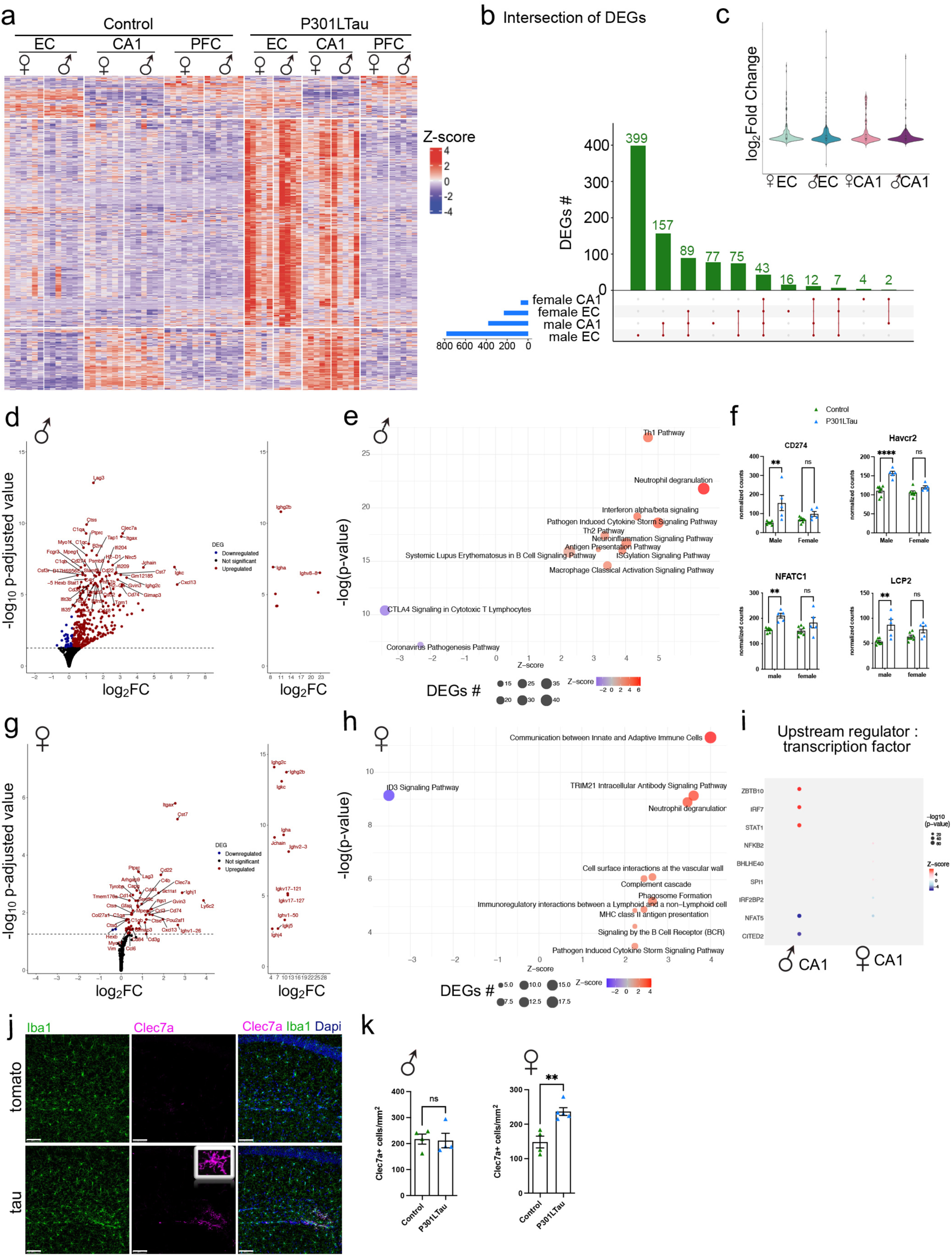
Tau injection alters EC and hippocampal gene pathways in a sex-specific manner. (**a**) Heatmap of differentially expressed EC and hippocampal genes (DEGs) from mice that tomato or tau injection (n = 5-7 mice per group, DESeq2, FDR<0.1). (**b**) Distribution of DEGs per brain region and sex, +/- tau injection contrats. (**c**) Log2 fold change of tau affected DEGs in females vs. males EC and CA1. (**d**) Volcano plot of tau DEGs in males (Log2FC>0.2; Padj<0.05). (**e**) Top-affected canonical pathways in tau CA1 males from IPA are represented. (**f**) Scatter plot representing DEGs from female and male mice, that are modulated by tau in males (FDR<0.1). (**g**) Volcano plot of tau DEGs in females (Log2FC>0.2; Padj<0.05). (**h**) Top-affected canonical pathways in tau CA1 females from IPA are represented. (**i**) Top CA1 tau Upstream regulators from IPA in both male and female mice. (**j**) representative images of iba1 (green), Clec7a (magenta) and dapi (blue) stainings in CA1 of tomato and tau mice (left). (**k**) Quantification of Clec7a+ microglial cells in CA1 of tomato and tau male and female mice (rigt). Graph indicates mean±s.e.m.

We then focused on the DEGs specifically affected in EC and CA1 in the comparison of Tau vs. tdTomato split by sex. At the EC level, we found 224 upregulated genes in females, 703 in males and 1 downregulated in females and 61 in males (**Suppl. Fig. 8a,c**; **Table 1**). Ingenuity Pathway Analysis showed very similar profiles in females and males in the comparison of Tau compared with tdTtomato groups. Indeed, among these pathways, we found in the top10 pathways, a common upregulation of neuroinflammation, Th1 and Th2 response, and phagosome pathways, and a common downregulation of CTLA4 signaling in cytotoxic T lymphocytes and ID3 signaling pathways (**Suppl. Fig. 8b,d).** Upstream regulator indicated a very similar profile of transcription factors induced by Tau in both males and females including the potential induction of ZBTB10 and the dowregulation of SIRT1, NFAT5, IRF2BP2 and CITED2. Potential induction of STAT1 was found to be specific to male EC, whereas IRF7 and IRF3 were found to be specific to female EC (**Suppl. Fig. 8e).**

At the CA1 level, a more dichotomic status was revealed. We found that male DEGs affected by tau in CA1 compared to tdTomato group, we found 350 upregulated and 20 downregulated genes (**Fig. 4d**, **Table 1**). Ingenuity Pathway Analysis showed in the top 10 pathways, an upregulation of Th1 and Th2 patways, of the neuroinflammation signaling pathway, and a downregulation of the CTLA4 signaling in cytotoxic T lymphocytes (**Fig. 4e, Table1**), also visible at the level of CD274 and HAVCR2 for the Th pathways and of NFATC1 and LCP2 gene counts for the neuroinflammation pathway (**Fig. 4i**).

In regards to female, DEGs affected by Tau propagation in comparison to tdTomato group included 64 upregulated and 2 downregulated genes (**Fig. 4g, Table1**). Among these, Ingenuity Pathway Analysis showed an upregulation of canonical pathways including in the top 10, communication between innate and adaptive immune cells as the highest one, induction of complement cascade and phagosome formation, and a downregulation of the canonical ID3 signaling pathway (**Fig. 4h**, **Table 1**).

Of note, in the CA1, several pathways appear still significantly affected in both sex, including complement cascade , phagosome formation, and T lymphocyte activation pathways. Among the differentially affected pathways between female and male, CTLA4, interferon alpha/beta signaling and neuroinflammation signaling pathways appear significantly different. Finally, upstream regulator analysis identified SIRT1 and CITED2 as downregulated upstream regulators and IR7 as an upregulated upstream regulator specifically in male CA1 compared to female CA1 (**Fig. 4i**). We then decided to explore the presence of T lymphocytes in the CA1 after Tau propagation. Using immunohistochemistry approach, we found a significant increase in CD4^+^ T lymphocytes in the CA1 of males and females after Tau propagation compared to tdTomato control group, with no effect on CD8^+^ T lymphocytes (**Suppl. Fig. 9**). Overall, these data indicate that there is a higher neuroinflammatory response in males than females in response to Tau propagation, and a global involvement of T lymphocytes in the response to Tau propagation.

Given that communication between immune cells was highlited directly in IPA terms or embedded in the neuroinflammatory response pathway, we decided to explore the effect of Tau propagation in CA1 with respect to the main cellular components we could isolate from the bulk RNAsequencing data. To do so, we analysed the expression levels of 1000 unique cellular genes from neuronal, astrocytic, oligodendrocytic and microglial origin [20]. When looking at the EC and CA1, our heatmaps visualization showed first that the two brain regions present a specific gene signature for neurons, astrocytes, microglia and oligodendrocytes. In addition, we found that only microglia present an altered gene expression signature in tau group compared to tdTomato, differences present in both males and females (**Suppl. Fig. 10**). Furthermore, we interrogated microglial genes, focusing on a set of genes representing the microglia in neurodegeneration (MGnD) signature [21]. We found that MGnD microglia were induced in response to Tau expression in the EC and also in the CA1, but to a lesser degree, of both males and females without a clear difference between the sexes at the heatmap visualization level (**Suppl. Fig. 11**). We finally conduced immunohistochemistry to quantify the density of microglia in the CA1 expressing Clec7a, a key gene of the MGnD signature associated with a high phagocytic activity of microglia, generally found in the vicinity of abnormal Tau accumulation or amyloid plaques [22]. Interestingly, Clec7a^+^ microglial density was found significantly higher in Tau group compared to tdTomato exclusively in females (**Fig. 4j-k**). In summary, the analyses from the bulk RNA-seq data demonstrate significant induction of neuroinflammation in EC and CA1 region in Tau mice, including Th1 and Th2 responces and phagosome pathways, and a common downregulation of CTLA4 signaling in cytotoxic T lymphocytes in EC and CA1 region of Tau male mice, while Tau female mice show induction of complement cascade and phagosome formation, and a downregulation of the canonical ID3 signaling pathway in the CA1 region. Immunohistochemical analysis show infiltration of CD4^+^ T cells in the CA1 region in both Tau male and female group, while female Tau mice show unique enrichment of activated MGnD microglial gene signature, and accumulation of Clec7a^+^ microglia in the CA1 region.

## Discussion

Here we used a circuit-defined model of early tau propagation to interrogate sex-specific consequences of entorhinal layer II to CA1 transfer; a pathway consistent with the earliest Braak-stage involvement of entorhinal cortex and hippocampal CA1 in Alzheimer’s disease. A key outcome is that tau propagation to CA1 (including along the proximal–distal axis) was comparable in males and females, yet functional impairment diverged sharply. Females developed deficits across multiple hippocampal-dependent domains (forced alternation, novel object recognition, fear acquisition, trace associative memory, and contextual fear memory), whereas males displayed a narrower impairment centered on trace associative memory. These data argue that sex differences emerge downstream of tau propagation, through distinct circuit and/or immune responses, rather than from differences in the anatomical burden of tau accumulation.

The pattern of behavioral impairment provides clues to which circuit operations are most vulnerable. Trace associative memory was disrupted in both sexes, consistent with temporoammonic pathway dysfunction affecting the timing-dependent association of discontinuous events [6, 8]. By contrast, the additional working memory, recognition, and contextual deficits in females point to broader degradation of CA1-dependent encoding and retrieval processes. Importantly, the female group showed increased locomotor measures in the Y-maze, but training-phase exploration in the object task was unchanged; together with the convergent deficits across paradigms, this supports a primary cognitive phenotype rather than reduced exploration or motivational confounds.

Electrophysiology of CA1 pyramidal neurons closely paralleled the behavioral divergence. In females, tau propagation produced a shift toward reduced excitability characterized by slower action potential kinetics, reduced spike output during depolarizing steps, and reduced spontaneous EPSC amplitude; changes expected to lower CA1 responsiveness and impair the fidelity of information transfer within hippocampal microcircuits. In males, intrinsic changes were modest (shorter action potential duration) and spontaneous EPSCs showed altered kinetics without a reduction in spike output, suggesting that the male phenotype may reflect altered synaptic timing and integration rather than a generalized hypoexcitability. The absence of detectable changes in dendritic spine density at this early time point indicates that functional alterations can precede overt structural synapse loss, highlighting an early window in which circuit physiology is disrupted before substantial remodeling.

Bulk transcriptomics revealed that tau propagation engages immune programs in both entorhinal cortex and CA1, consistent with emerging views that tau-associated network dysfunction engages both innate and adaptive immune components [23], but with sex-modulated signatures. A striking observation is that males exhibited a larger number of differentially expressed genes, with prominent Th1/Th2 and neuroinflammatory pathway enrichment and CTLA4-associated signaling changes, while females showed fewer DEGs yet a more CA1-focused enrichment of complement and phagosome pathways and communication between innate and adaptive immune cells. This apparent dissociation, more transcriptional change in males but broader physiological/behavioral impairment in females, suggests that the phenotype and cellular targets of the immune response, rather than its overall magnitude, may determine functional outcome. Bulk RNA-seq integrates shifts in cell states and cell proportions; thus, differences could reflect distinct activation states of microglia and infiltrating immune populations that have different functional consequences on synapses and excitability.

Two findings support a model in which complement-linked microglial programs may have heightened functional impact in females. First, females showed enrichment of complement–phagosome pathways in CA1, which are positioned to drive synaptic weakening through tagging and microglial engulfment. Second, only females exhibited increased density of Clec7a^+^ microglia in CA1, a marker associated with MGnD/DAM-like states and elevated phagocytic potential. Together with the reduced sEPSC amplitude and reduced firing capacity in females, these results are consistent with preferential disruption of excitatory synaptic function in CA1. Complement system, especially C1q and C3aR, plays a key role for the progression of synaptic loss in PS19 mice [24, 25]. Strong MGnD/DAM induction in Tau female mice may attribute to the synaptic impairment in the CA1 region. In contrast, male-biased Th-related and CTLA4-linked signatures may reflect differences in adaptive immune regulation and cytokine milieu that modulate synaptic kinetics and timing-dependent plasticity (impacting trace memory) without producing the broader reduction in excitability seen in females. Notably, CD4^+^ T-cell infiltration was present in both sexes, indicating a shared adaptive immune component; whether the activation state, antigen specificity, or effector programs differ by sex remains an important open question.

This study has a few limitations. Future work should resolve causality and cellular specificity by integrating single-nucleus or spatial transcriptomics with targeted perturbations. Testing whether complement blockade, microglial phagocytosis inhibition, or T-cell depletion differentially rescues CA1 physiology and behavior in females versus males would directly evaluate the proposed mechanisms. Longitudinal analyses will also be needed to determine whether early functional changes progress to structural synapse loss and whether immune signatures evolve over time in a sex-dependent manner. Overall, our findings support a framework in which early entorhinal-to-CA1 tau propagation triggers sex-specific neuroimmune programs that shape CA1 neuronal dysfunction and cognitive vulnerability, providing a tractable model to dissect early-stage mechanisms relevant to sex differences in AD.

## Acknowledgment

We would like to thank Dr. Susumu Tonegawa for Wfs1-Cre mouse model and members of Molecular Neurotherapeutics Laboratory at Boston University School of Medicine and Mayo Clinic Florida.

## Funding

This work was funded in part by NIH R01 AG066429 (T.I.), NIH R01 AG072719 (T.I.), NIH R01 AG067763 (T.I.), NIH RF1 AG054199 (T.I.), NIH R01 AG054672 (T.I.), Cure Alzheimer’s Fund (T.I., S.I.), Alzheimer’s Association Zenith Fellows Award ZEN-26-1436830 (T.I.), the INRAE DIGIT-BIO metaprogram DINAMIC grant (JCD, CM, AR), the Région Nouvelle-Aquitaine CHESS Exomarquage 13059720-13062120 (JCD, CM).

## Author contributions

Conceptualization, T.I.; Methodology, J.C.D., D.P., C.M.,T.I., S.I., J.I.L., M.M.; Investigation, JC.D., S.V.K, D.P., H.V.L, C.M.; Writing – Original Draft, J.C.D.,S.I., T.I.; Writing – Review & Editing, all authors; Validation, ; Formal Analysis: J.C.D., S.V.K., D.P., J.I.L., C.M., A.R. H.V.L.; Funding Acquisition, T.I., S.I.,J.C.D.; Supervision, T.I, J.I.L., S.I.

## Author information

To be added upon request

## Conflict of Interest

The following authors have competing financial interests: T.I. is on SAB of AAVINE and consults for Takeda and Otsuka Pharma.

---

**Supplemental Fig. 1.**
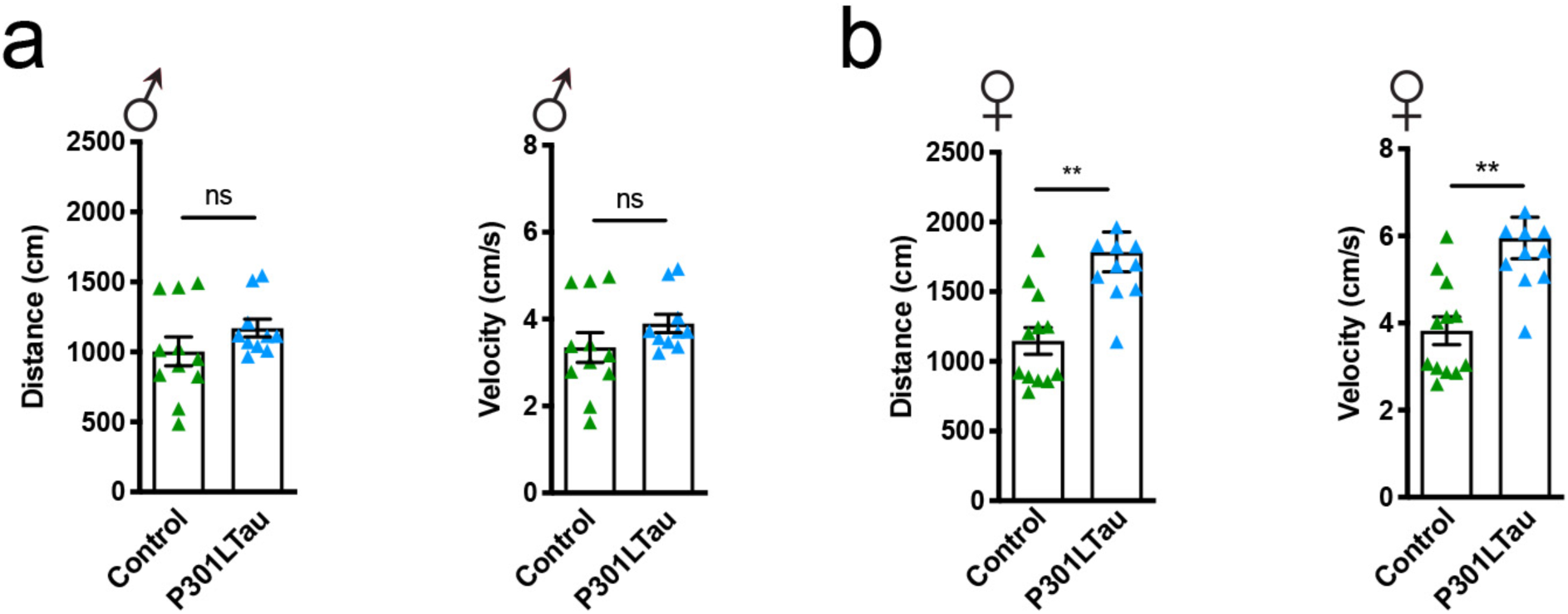
Motricity evaluation. **(a)** Total distance explored (left) and mean velocity (right) in the Y-maze for male during the test session showing no difference between tomato and tau groups. **(b)** Total distance explored (left) and mean velocity (right) in the novel object recognition test for female during the test session showing no difference between tomato and tau groups. Graph indicates mean±s.e.m.

**Supplemental Fig. 2.**
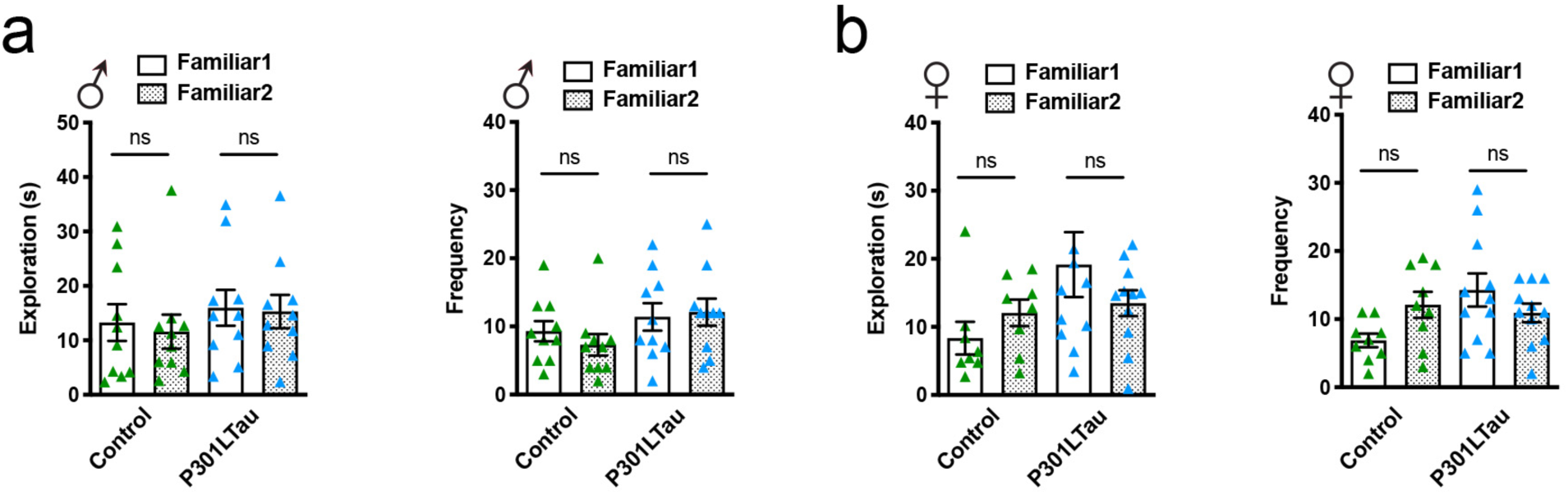
Novel object recognition training phase. **(a)** Total exploration duration (left) and frequency (right) in the novel object recognition test for male during the training session showing no difference between tomato and tau groups. **(b)** Total exploration duration (left) and frequency (right) in the novel object recognition test for female during the training session showing no difference between tomato and tau groups. Graph indicates mean±s.e.m.

**Supplemental Fig. 3.**
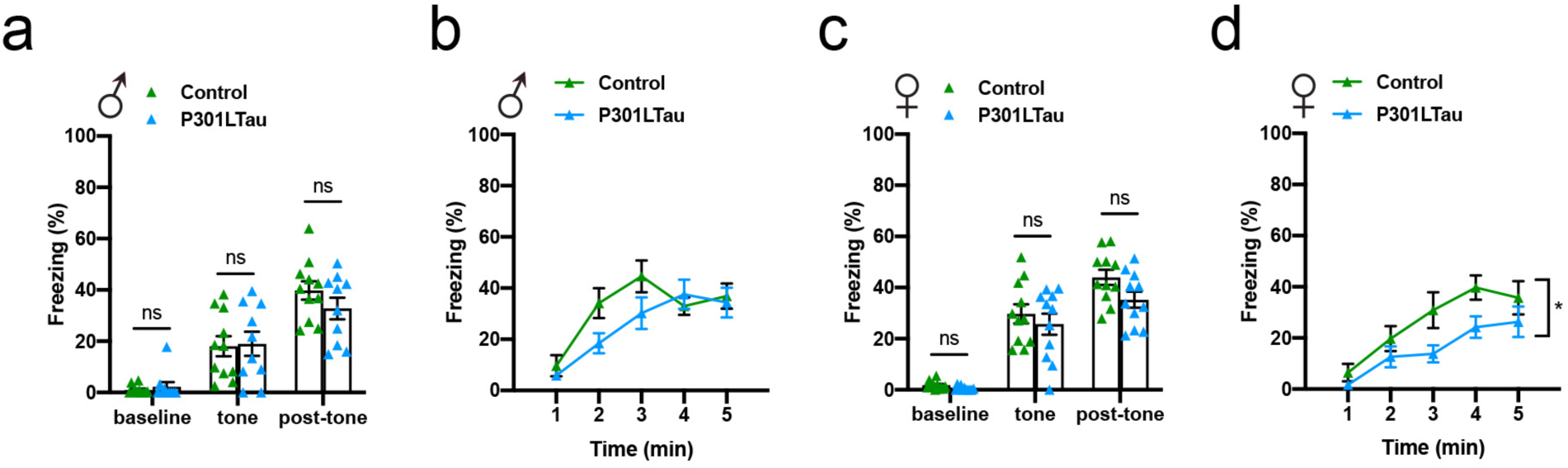
Fear conditioning additional informations. **(a)** Average freezing time during baseline, tone and post-tone periods measured during the training phase of the fear conditioning protocol on day 1 in males, showing no differences between tomato and tau groups. (**b**) Freezing time during contextual-memory testing observed on day 3 in males, showing no difference among the groups. **(c)** Average freezing time during baseline, tone and post-tone periods measured during the training phase of the fear conditioning protocol on day 1 in females, showing no differences between tomato and tau groups. (**d**) Freezing time during contextual-memory testing observed on day 3 in females, showing a significant difference between tomato and tau groups. Graph indicates mean±s.e.m.

**Supplemental Fig. 4.**
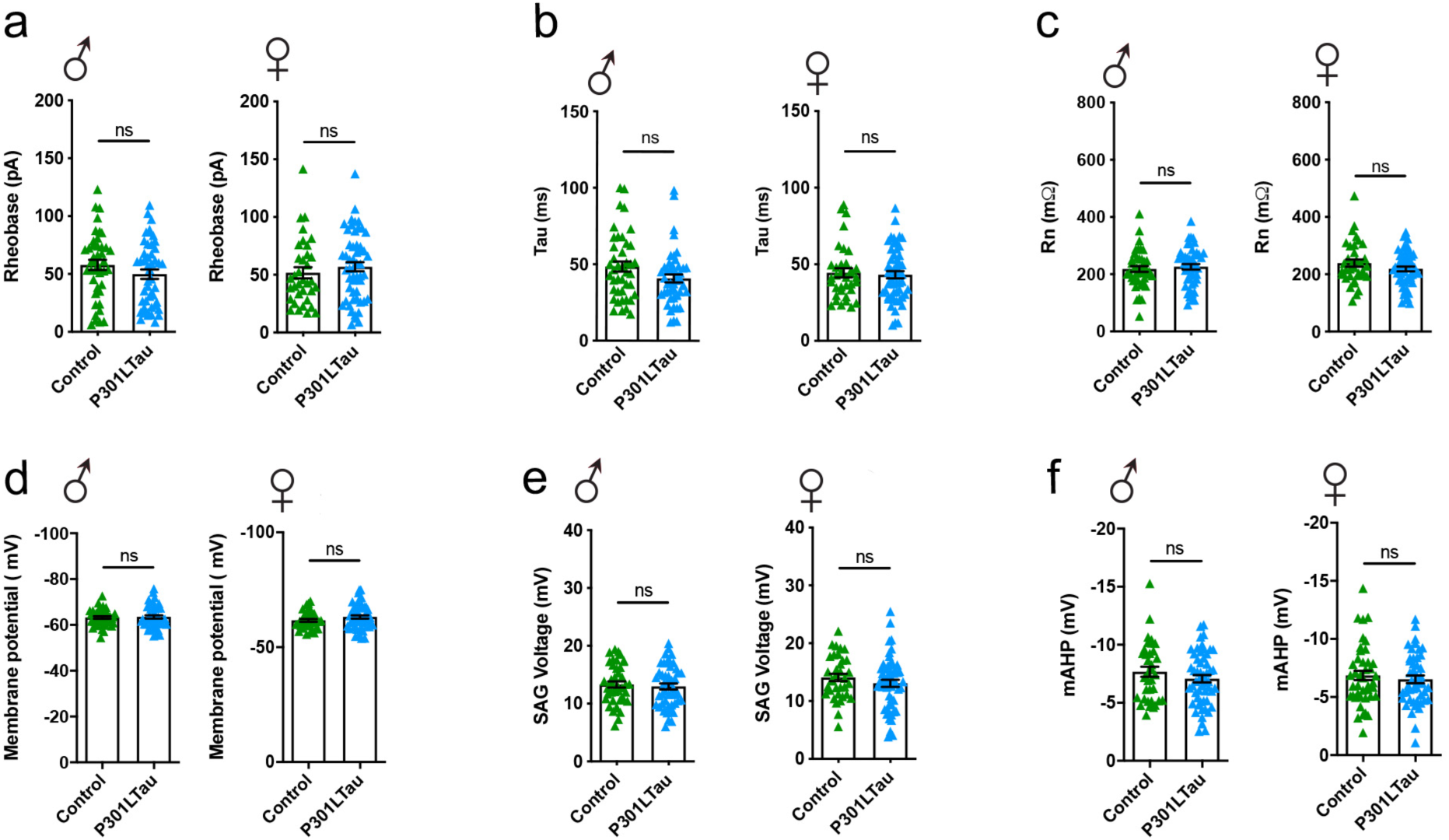
**Instrinsic additional properties of pyramidal neurons of CA1 in both sexes and tomato and tau groups**. (**a**) rheobase; (**b**) Tau; (**c**) Resistance; (**d**) Membrane resting potential; (**e**) SAG voltage; (**f**) mAHP; showing no difference between the groups. Graph indicates mean±s.e.m.

**Supplemental Fig. 5.**
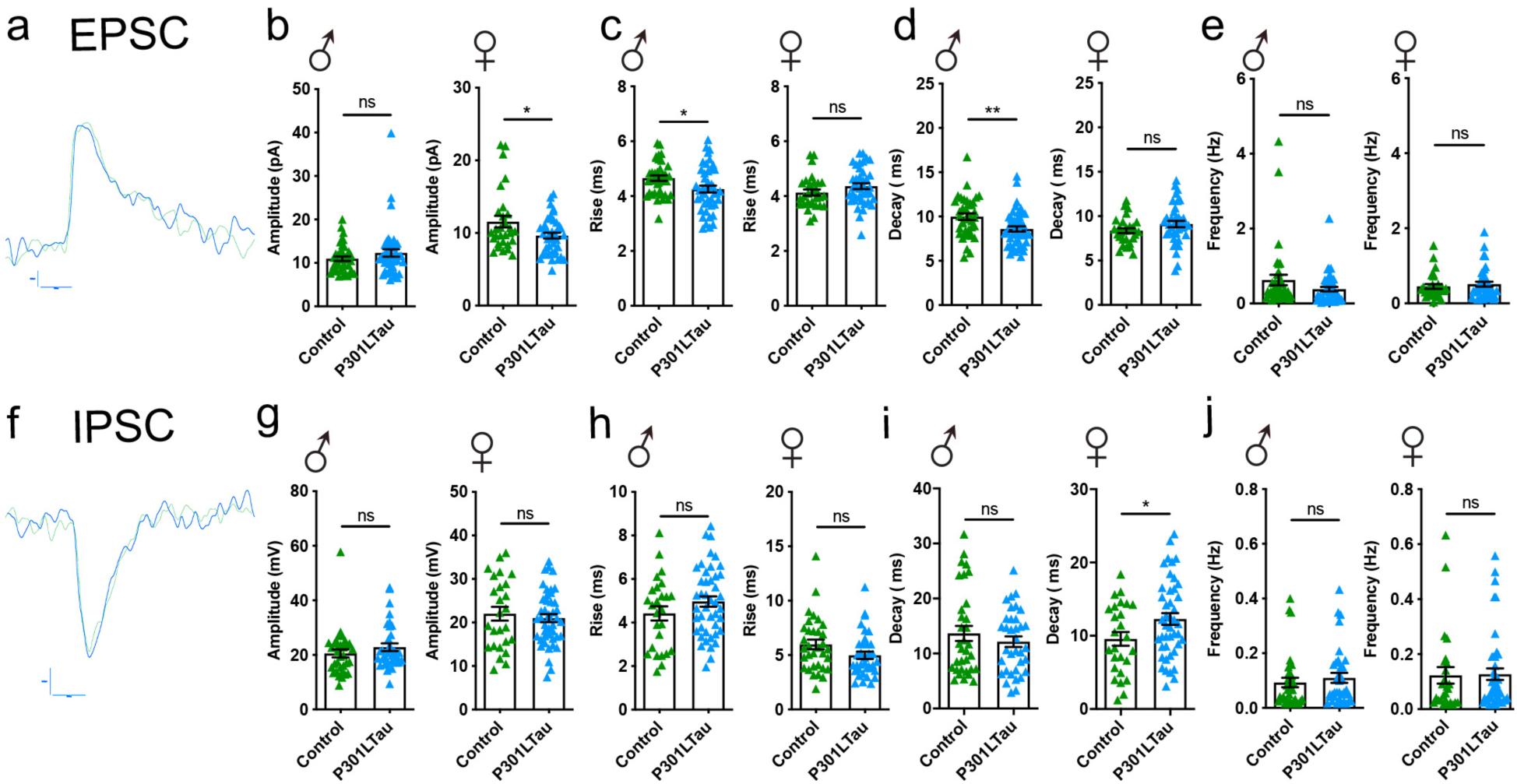
Excitatory and inhibitory post synaptic current parameters. **(a)** EPSC representative trace. **(b)** EPSC amplitude. **(c)** EPSC rise. **(d)** EPSC decay. **(e)** EPSC frequency. **(f)** IPSC representative trace. **(g)** IPSC amplitude. **(h)** IPSC rise. **(i)** IPSC decay. **(j)** IPSC frequency. Graph indicates mean±s.e.m.

**Supplemental Fig. 6.**
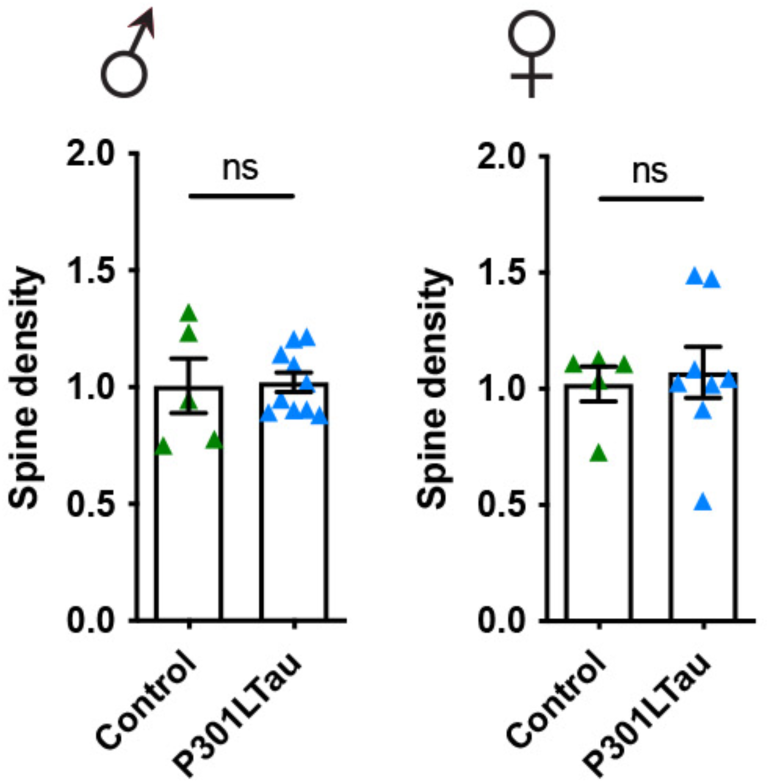
Spine neuronal analysis. Spine density measured from recorded and biocytin-filed CA1 pyramidal neurons from both sexes and tomato and Tau groups, showing no difference among the groups. Graph indicates mean±s.e.m.

**Supplemental Fig. 7.**
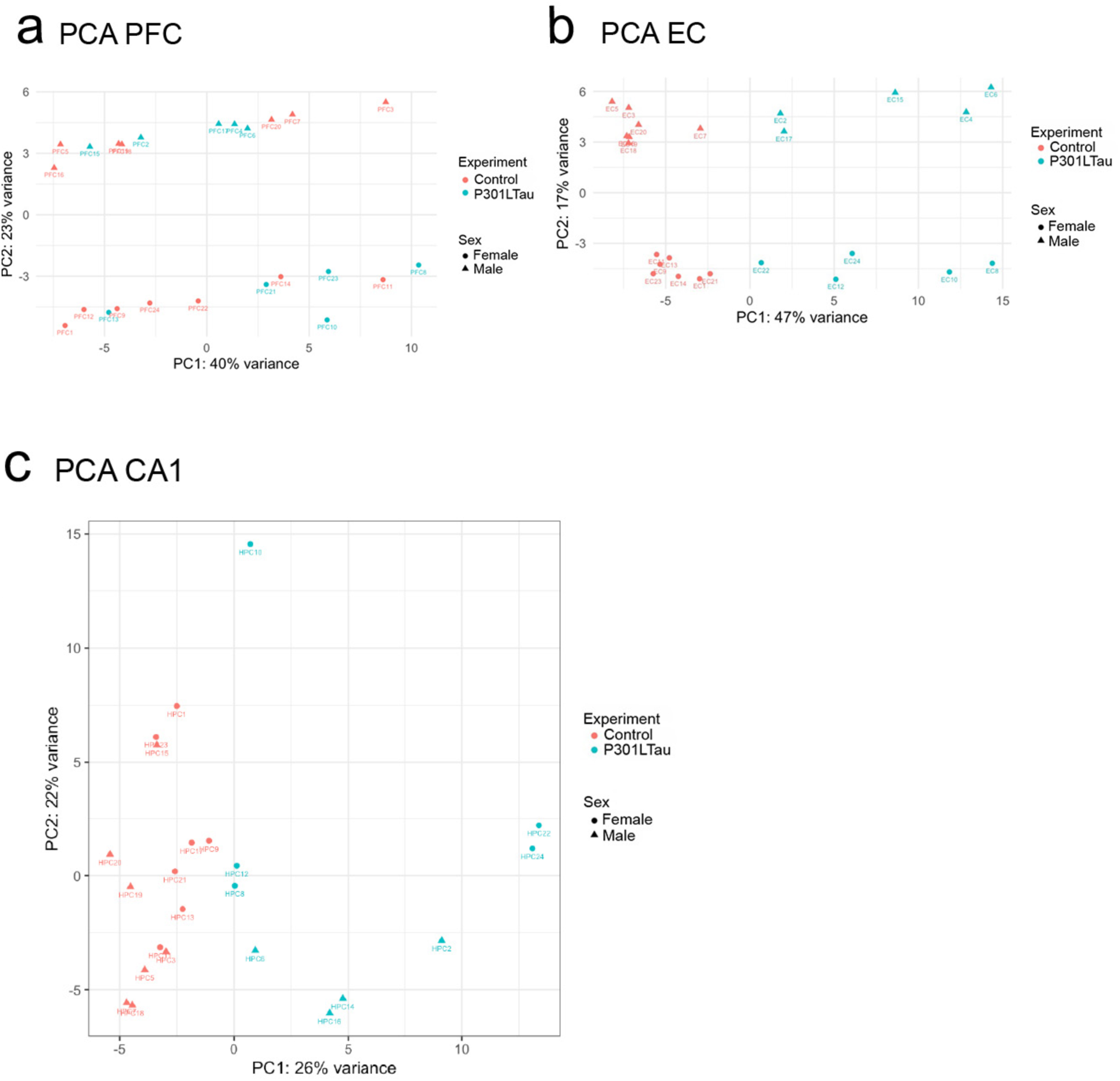
Principal component analysis diagrams of DEGs from both sexes and tomato and Tau groups: in prefrontal cortex (PFC) **(a)**; enthorinal cortex (EC) **(b)**; and hippocampal CA1 subregion **(c)**, showing only a sexual separation in PFC, a treatment and sex separation in EC and CA1.

**Supplemental Fig. 8.**
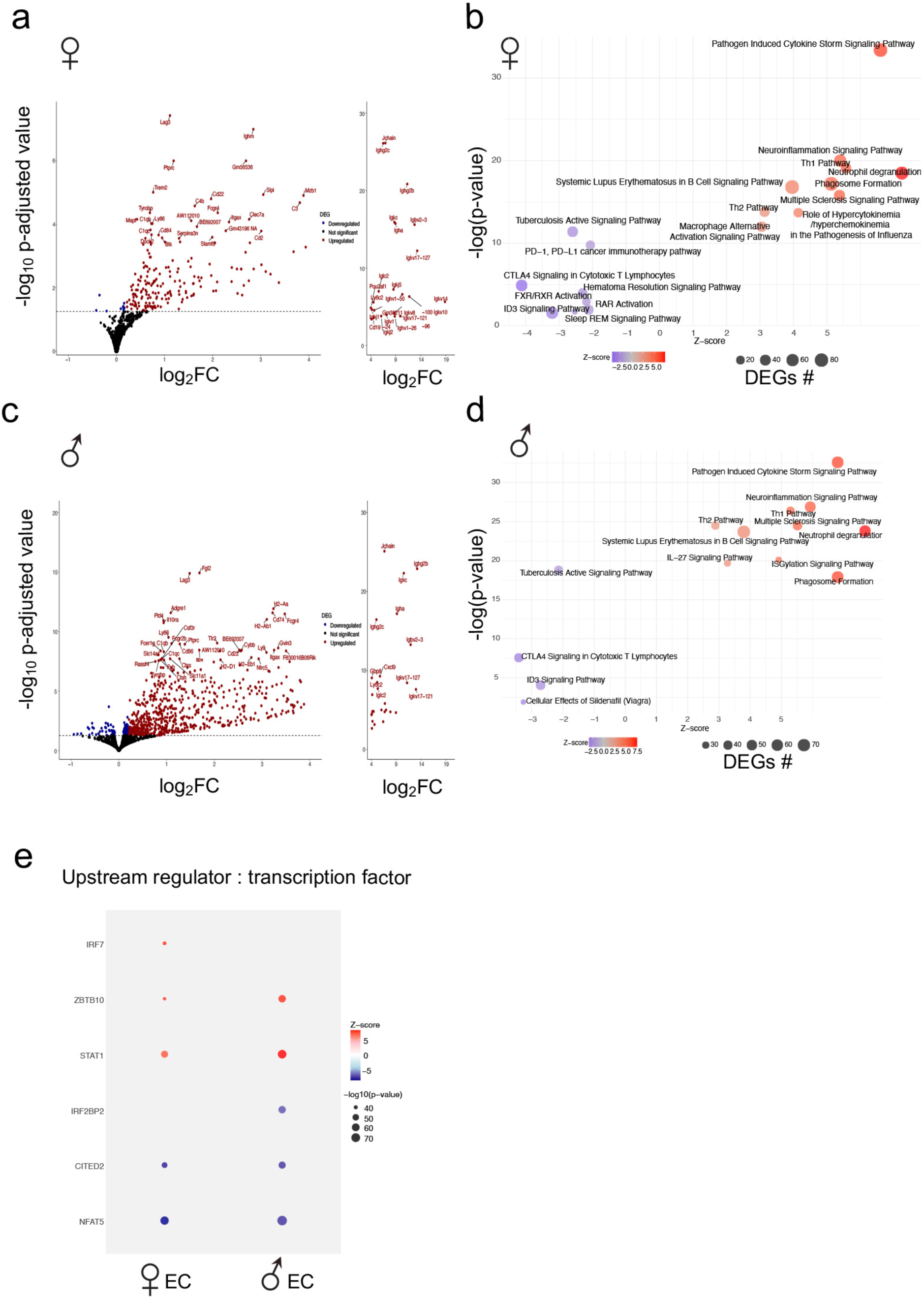
RNAsequencing analysis of EC region. (**a**) Volcano plot of tau DEGs in females. (**b**) Top-affected canonical pathways in tau EC females from IPA are represented. (**c**) Volcano plot of tau DEGs in males. (**d**) Top-affected canonical pathways in tau EC males from IPA are represented. (**e**) Top EC tau Upstream regulators from IPA in both male and female mice.

**Supplemental Fig. 9.**
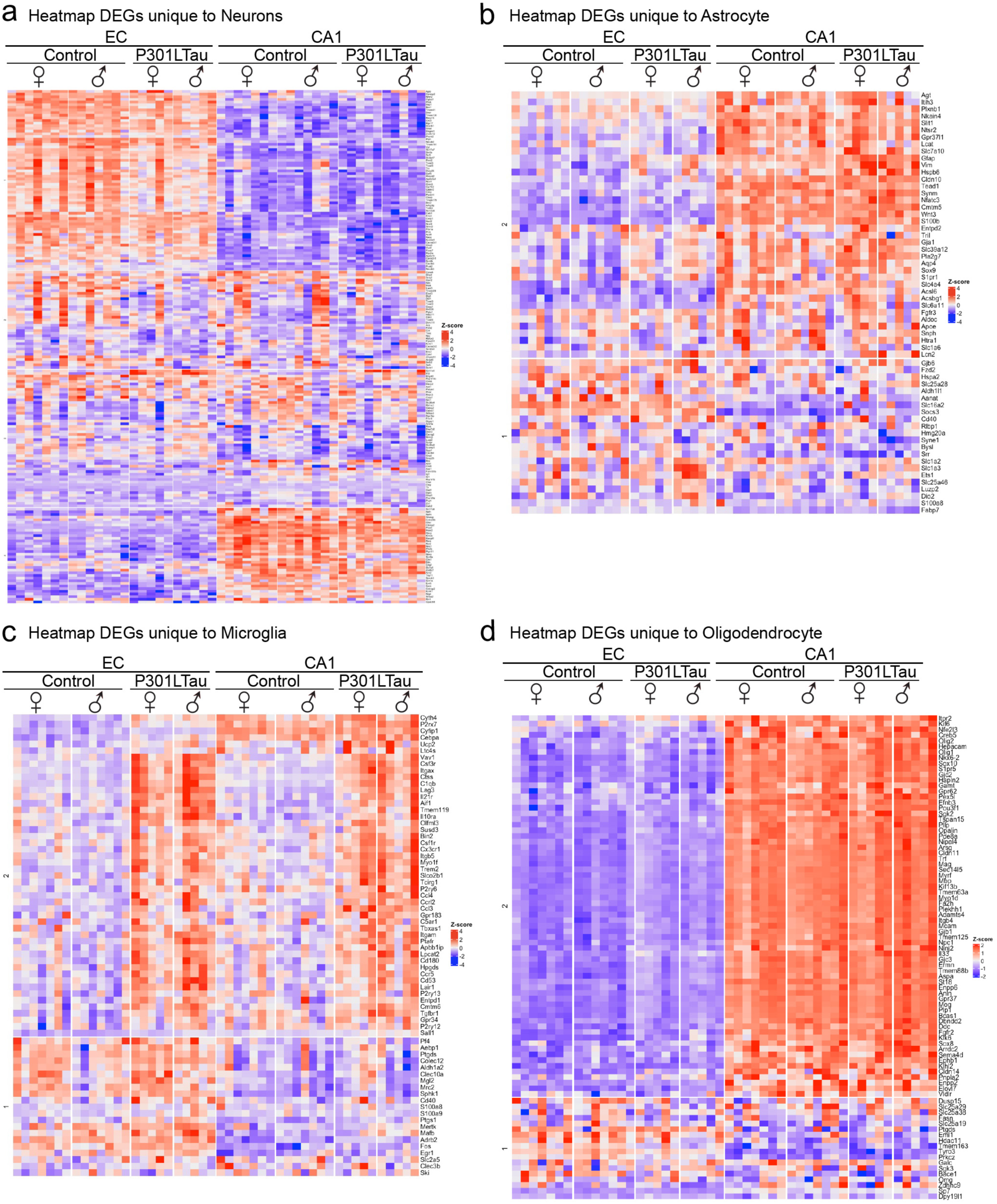
Heatmaps generated using a database of 1000 unique genes per cell type. **(a)** heatmaps of the detected genes out of 1000 unique neuronal genes from both sexes and tomato and Tau groups. **(b)** heatmaps of the detected genes out of 1000 unique astrocytic genes from both sexes and tomato and Tau groups. **(c)** heatmaps of the detected genes out of 1000 unique microglial genes from both sexes and tomato and Tau groups. **(d)** heatmaps of the detected genes out of 1000 unique oligodendrocytic genes from both sexes and tomato and Tau groups.

**Supplemental Fig. 10.**
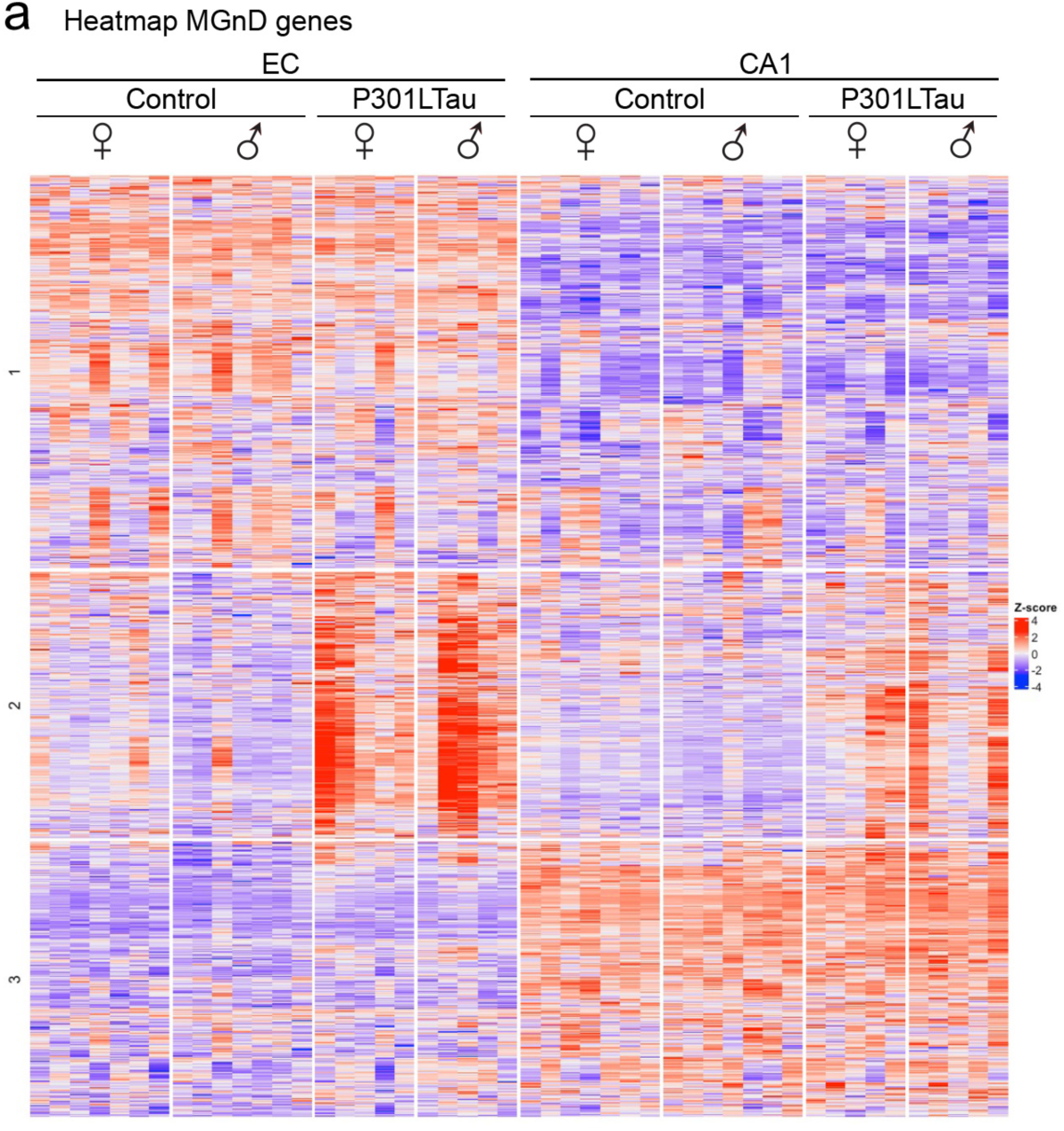
Microglial MGnD signature evaluation. **(a)** heatmaps of the detected MGnD signature genes from both sexes and tomato and Tau groups.

**Supplemental Fig. 11.**
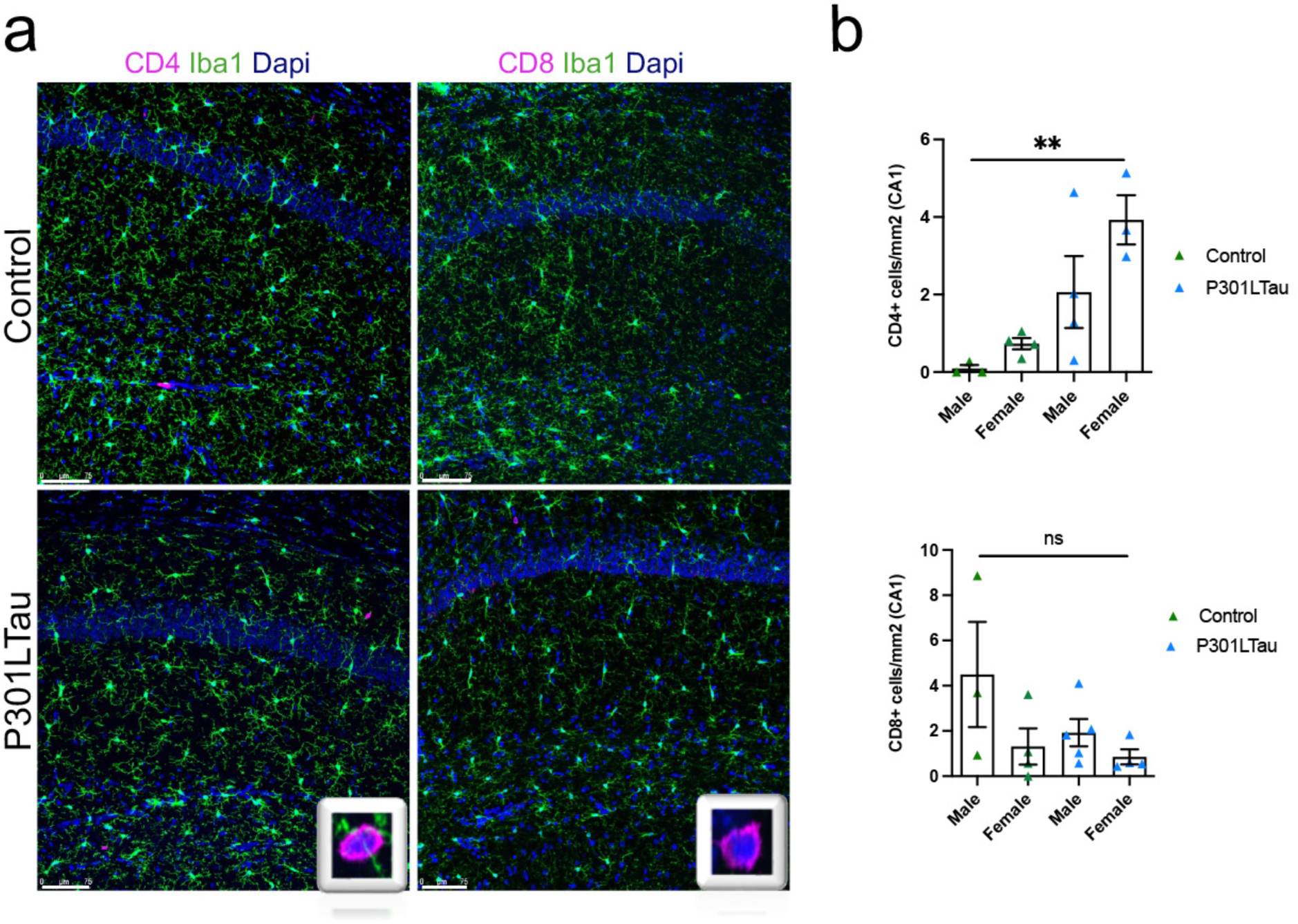
T lymphocyte infiltration evaluation. **(a)** representative images of iba1 (green), CD4 or CD8 (magenta) and dapi (blue) stainings in CA1 of tomato and tau mice. **(b)** Quantification of CD4^+^ T lymphocytes (top) or CD8^+^ T lymphocytes (bottom) in CA1 of tomato and tau male and female mice. Graph indicates mean ± s.e.m.

